# A Bioinformatically Identified 5-SNP Streak in the ARMS2 Promoter Region Links to AMD Through Altered c-ABL Binding and Transcriptional Regulation

**DOI:** 10.1101/2025.10.27.684937

**Authors:** Ping-Wu Zhang, Sheng Liu, Weifeng Li, Laura Fan, Sean Li, Zi-He Wan, Junyeop Ahn, Cynthia A. Berlinicke, Shannath L. Merbs, Donald J. Zack

## Abstract

Genome-wide association studies (GWAS) have identified the 10q26 locus as a major genetic risk factor for age-related macular degeneration (AMD), yet the causal variants and underlying mechanisms remain unresolved. Fine-mapping studies localized the AMD-associated region to a 5,196 bp interval near the ARMS2–HTRA1 locus, excluding both HTRA1 and the ARMS2 3′ indel variant as primary risk determinants. Using an unbiased bioinformatic approach based on protein–DNA interaction potential, we identified a polymorphic 5C/5A SNP streak within this interval predicted to alter binding of the non-receptor tyrosine kinase c-ABL. Electrophoretic mobility shift assays and chromatin immunoprecipitation–qPCR confirmed direct c-ABL binding to this regulatory element. The 5-SNP streak is in complete linkage disequilibrium with rs10490924 (A69S) and showed strong association with both wet AMD (OR = 3.37) and dry AMD (OR = 3.80) in 566 human DNA samples. Luciferase reporter assays demonstrated reduced transcriptional activity of the risk 5A haplotype in the presence of c-ABL. Consistent with these findings, ARMS2 mRNA expression was significantly increased (1.73-fold) in human RPE/choroid tissues carrying the risk 5A haplotype, whereas HTRA1 expression was unchanged. These results identify a functional noncoding regulatory element at 10q26 and implicate altered c-ABL–mediated regulation of ARMS2 in AMD pathogenesis.

## Introduction

Age-related macular degeneration (AMD), a leading cause of irreversible blindness in elderly populations, is characterized by progressive degeneration of photoreceptor (PR) and the retinal pigment epithelial (RPE) cells in the macula region. Clinically, AMD manifests as two major forms: wet (neovascular) AMD, which accounts for 10–15% of cases and is more severe, and dry (atrophic) AMD, which represents 85–90% of cases. By 2040, the global prevalence of AMD is projected to rise to nearly 288 million individuals [1]. While anti-VEGF therapies have transformed the management of wet AMD, effective treatments for dry AMD remain limited. Recently developed complement-targeting therapies (C3 and C5 inhibitors) offer promise, but their efficacy remains controversial [2].

AMD arises from a complex interplay of environmental and genetic factors. Aging, smoking, and diet are established non-genetic risk factors, while genetic studies have consistently highlighted two susceptibility loci: 1q31 (CFH) and 10q26 (*ARMS2*–*HTRA1*) [3–6]. The missense SNP rs10490924 (c.205G>*T*, p.A69S, T is the risk allele) in *ARMS2* and rs11200638 in the *HTRA1* promoter have been studied extensively, but strong linkage disequilibrium across the region complicates determination of causality [7–9]. Despite strong associations across diverse populations, the precise causative variants and pathogenic mechanisms at the 10q26 locus remain unresolved [10,11]. Copy number variation analyses have excluded large structural variants, suggesting that functional contributions most likely arise from SNPs or small indels [12,13].

Functional studies have produced conflicting results. *ARMS2*, a primate-specific gene, has no clear phenotype in transgenic mice [14,15]. *HTRA1*-knockout mice also show no retinal changes at 12 months [16], though overexpression of *HTRA1* has been linked to pathology in models of polypoidal choroidal vasculopathy and zebrafish photoreceptor degeneration [17,18]. Additional complexity arises from isoform-specific regulation; for example, *ARMS2*-B is expressed at levels nearly 30-fold higher than *ARMS2*-A [19]. Studies of *ARMS2* and *HTRA1* expression in AMD patient tissues have further yielded inconsistent findings [20].

Refinement of the 10q26 risk locus has provided new clues. Fritsche et al. mapped the association to a 30 kb interval upstream of *ARMS2*, excluding the *ARMS2* p.R38X variant [21]. Grassmann et al., analyzing 16,144 AMD cases and 17,832 controls, narrowed the risk haplotype to a 5,196 bp segment encompassing the *ARMS2* promoter and first two exons, while excluding the *HTRA1* promoter and *ARMS2* 3′ indel variant [22]. These results suggest the presence of asyet unidentified regulatory elements in this region.

Taken together, the evidence points toward noncoding regulatory variants as drivers of risk at the 10q26 locus. To address this gap, we combined genomics, bioinformatics, and functional assays to systematically investigate variants within the 5,196 bp *ARMS2* promoter region, with a particular focus on SNPs that may modulate protein–DNA interactions. Our goal was to uncover novel regulatory elements that contribute to AMD pathogenesis.

### Ethics statement

All aspects of this study were conducted in accordance with the principles of the Declaration of Helsinki, with informed consent being obtained from all participants. This project was approved by the Johns Hopkins University School of Medicine Institutional Review Board (IRB).

## Material and Methods

### Bioinformatic prediction of SNP–protein interactions

Within the 5,196 bp region (chr10:122,453,933–122,454,174, GRCh38), 27 common SNPs (minor allele frequency >0.1) were analyzed. For each SNP, 30 bp flanking sequences were scanned using TFBSTools [23] and 6,088 motifs curated from published studies and the MotifDB database [24]. Motif enrichment was filtered using a Benjamini–Hochberg adjusted p-value <1 × 10□^3^. Transcription Factor Affinity Prediction (TRAP) [25-27] was then applied to calculate motif affinities for both reference and alternate alleles. Only motifs with affinity p-values <1 × 10□□ and significant allele-dependent changes were retained.

### Human AMD eye samples

Post-mortem eyes were obtained from the National Disease Research Interchange (NDRI, Philadelphia, USA) and transported on wet ice. RPE/Choroid tissues were isolated within 48 hours of death. All tissue handling and processing procedures complied with the ARVO Best Practices for Using Human Eye Tissue in Research (Nov 2021). 298 Blood DNA samples ware mentioned in our previous paper [28]. Others were further collected in our lab.

### RNA extraction from cultured cells

Total RNA was extracted from ∼50 mg of retinal tissue using TRIzol (Thermo Fisher Scientific, Invitrogen, USA) and further purified the PureLink RNA Mini Kit (Invitrogen, USA) with on-column DNase digestion, following the manufacturer’s protocol. Samples were processed with chloroform extraction and ethanol precipitation. RNA was purified on spin cartridges and eluted in RNase-free water. RNA concentration and purity were assessed with a NanoDrop spectrophotometer (Thermo Fisher Scientific, USA), and only samples with an A260/280 ratio between 1.7 and 2.1 were used for downstream assays.

### Reverse transcription and quantitative -PCR (qPCR)

Complementary DNA (cDNA) was synthesized from 500 ng of total RNA in a 20 μL reaction volume using either the High-Capacity cDNA Reverse Transcription Kit (Applied Biosystems) or the iScript™ cDNA Synthesis Kit (Bio-Rad), according to the manufacturers’ instructions.

Quantitative PCR (qPCR) was performed using iTaq Universal SYBR Green Supermix (Bio-Rad) on a Bio-Rad CFX384 Real-Time PCR system. All reactions were performed in biological triplicates. qPCR reactions were set up in a final volume of 5 μL per well, containing 2.5 μL of 2× SYBR Green Master Mix, 0.25 μL each of forward and reverse primers (final concentration 250 nM), 1.5 μL of 3-fold diluted cDNA template (or 1.5 μL of undiluted cDNA for *ARMS2*), and 0.75 μL of nuclease-free water. Reactions were dispensed into 384-well optical plates, sealed with optical adhesive film, and briefly centrifuged to remove air bubbles.

Amplification was performed on a 384-well real-time PCR system using the following cycling conditions: an initial denaturation at 95 °C for 30 s, followed by 40 cycles of denaturation at 95 °C for 5 s and annealing/extension at 60 °C for 25 s. A melt curve analysis was conducted at the end of amplification to confirm primer specificity. Gene expression levels were normalized to the geometric mean of two housekeeping genes, GAPDH and ACTB. Primer sequences were as follows: c-ABL forward, AGAGCTGCAGAGCACAGAGA; c-ABL reverse, CTCTTTTCGAGGGAGCAATG; GAPDH forward, CTGACTTCAACAGCGACACC; GAPDH reverse, TAGCCAAATTCGTTGTCATACC; ACTB forward, GCCTCGCCTTTGCCGAT; ACTB reverse, CGCGGCGATATCATCATCCA; ARMS2 forward, CTCTGCGAGAGTCTGTGCTG; and ARMS2 reverse, ACAGGAGGGAATGCAGTGAC.

### Genomic DNA extraction

Genomic DNA from peripheral blood was extracted from whole blood using the Puregene Blood Core Kit C (Qiagen, USA). Genomic DNA (gDNA) from human eye tissues was isolated using the Quick-DNA MicroPrep Kit (Zymo Research, USA). Human eye tissues (5–10 mg, fresh or frozen) were homogenized in 500 µL Genomic Lysis Buffer (Quick-DNA Microprep Kit, Zymo, USA and SCILOGEX D160 homogenizer) and digested with proteinase K (final concentration 1 mg/mL; Invitrogen, USA) at 60 °C for 2–3 h. Lysates were centrifuged at 10,000 × g for 5 min, and the supernatant was applied to Zymo-Spin IC columns. Wash steps were performed sequentially with DNA Pre-Wash Buffer (200 µL) and gDNA Wash Buffer (500 µL), each followed by centrifugation at 10,000 × g for 1 min. DNA was eluted in 50 µL DNA elution buffer after 5 min incubation at room temperature and recovered by centrifugation at 10,000 × g for 1 min.

### Sanger sequencing

DNA sequencing was performed at the Johns Hopkins Genetics Resources Core Facility using an Applied Biosystems 3730xl DNA Analyzer (Thermo Fisher Scientific, USA).

### Electrophoretic mobility shift assay (EMSA)

A PCR fragment of 220 bp covering the 5-SNP streak was used to test the binding between c-ABL protein and the 5-SNP streak using a published method [29]. The 220 bp PCR fragment was amplified using primers: forward 5′-TGAGGCCACAATAAGCTGTG-3′ and reverse 5′-GGTAAGCATTCGGGAAATCTG-3′. PCR cycling conditions were: 98 °C for 1 min, followed by 30 cycles of 98 °C for 10 s, 61 °C for 10 s, and 72 °C for 20 s. Binding reactions contained 1.91 × 10□□ M DNA (∼100 ng) and increasing concentrations of recombinant human c-ABL protein (0.08–1.32 × 10□□ M; ab136358, Abcam, UK) in binding buffer (10 mM Tris, pH 7.6, 50 mM KCl, 1 mM DTT, 10 μg/mL BSA). Reactions were incubated on ice for 30 min and resolved on 1.5% agarose gels at 100 V for 50 min.

### Genotyping of SNP rs10490924 and the 5-SNP block

SNP rs10490924 was genotyped by PCR amplification of a 222-bp fragment, followed by restriction enzyme digestion with PvuII and analysis by agarose gel electrophoresis. Wild-type homozygous G/G alleles yielded two fragments (139 bp and 83 bp), homozygous T/T alleles remained undigested (222 bp), and heterozygous G/T samples displayed all three bands. PCR amplification was performed using the following primers: forward 5′-GCAAGTCTGTCCTCCTCGGTG-3′ and reverse 5′-GTGCTGAGGCTGCTGGAACC-3′.

The 5-SNP streak was genotyped by PCR amplification followed by Sanger sequencing using the same primer pair employed for EMSA, generating a 220-bp amplicon. Sequencing was performed using the reverse primer 5′-GGTAAGCATTCGGGAAATCTG-3′.

### Cell cultures

ARPE-19 and HEK293T cells were maintained at 37 °C in a humidified atmosphere (5% COL, 95% air) in Dulbecco’s modified Eagle’s medium (DMEM; 4.5 g/L glucose, L-glutamine, pyruvate) supplemented with 1% heat-inactivated fetal calf serum (FBS, Sigma Aldrich, USA) and 1% penicillin/streptomycin (Sigma Aldrich, USA). HEK293T cells were cultured in DMEM/10% FBS and 1% penicillin/streptomycin.

### Chromatin immunoprecipitation and quantitative PCR (ChIP-qPCR) assay

ARPE-19 Cells were cultured for 14 days and cross-linked with 1% formaldehyde in DMEM/10% FBS for 30 min at room temperature, quenched with 0.125 M glycine, and washed three times with ice-cold PBS containing 0.5 mM PMSF. Nuclei were isolated, resuspended in sonication buffer, and sonicated (VibraCell Sonicator, USA) 12 times for 10 s each. Lysates were pre-cleared with Dynabeads Protein A beads (ThermoFisher Sci, USA) and incubated overnight with 5 µg of anti-c-ABL antibody (8E9, Catalog sc-56887, Sant Cruz Bio, USA) followed by Protein A Sepharose capture. Immunoprecipitates were washed sequentially with buffers A, B, and TE, eluted in 200 µL elution buffer, and DNA was purified by phenol/chloroform extraction and ethanol precipitation with glycogen and sodium acetate. Enriched DNA was analyzed by quantitative PCR. These primers were used: forward 5′-TGAGGCCACAATAAGCTGTG-3′ and reverse 5′-GGTAAGCATTCGGGAAATCTG-3′.

### Luciferase reporter constructs

A 2.3-kb fragment upstream of ARMS2 was amplified from non-AMD genomic DNA (rs10490924, GG genotype) and cloned into the pGL3-Basic vector (Promega) using KpnI and XhoI restriction sites. A 100-bp subfragment flanking the 5-SNP streak within the 2.3-kb region was then replaced with the corresponding fragment containing either the wild-type sequence (AACAACAACAACAAAAAAACAA) or the mutant sequence (AAAAAAAAAAAAAAAAAAAAAACAA) using Gibson Assembly (New England Biolabs, USA). The integrity of the resulting 2.3-kb constructs harboring either the 5C or 5A variant was confirmed by Sanger sequencing.

### Luciferase Reporter Assay

HEK293 and ARPE-19 cells were cultured for 2 days and 10 days, respectively, before transfection. Cells were seeded in 24-well plates and transfected at ∼70–80% confluency with firefly luciferase reporter constructs using Lipofectamine 3000 (Thermo Fisher Scientific), according to the manufacturer’s instructions. For each well, cells were co-transfected with the firefly luciferase reporter plasmid and a Renilla luciferase plasmid (pRL-TK) as an internal control to normalize for transfection efficiency. The total amount of DNA was kept constant across conditions by supplementing with empty vector.

At 48 h post-transfection, cells were lysed using Passive Lysis Buffer (Promega). Luciferase activity was measured using the Dual-Luciferase® Reporter Assay System (Promega) following the manufacturer’s protocol. Firefly and Renilla luminescence signals were detected sequentially using a microplate luminometer. Firefly luciferase activity was normalized to Renilla luciferase activity for each sample. Data were collected from at least three independent biological replicates, each performed in technical triplicates.

### Statistical analysis

All data were analyzed using GraphPad Prism (GraphPad Software, USA) and Microsoft Excel. Statistical significance was evaluated using Student’s *t*-test or chi-square test, as appropriate. *P* < 0.05 was considered significant. Data are presented as mean ± SEM from three biological replicates. * Indicates P < 0.05, ** indicates P < 0.01, and *** indicates P < 0.001. ns, not significant.

## Results

### Bioinformatic screening identifies c-ABL as a candidate regulator within the 5,196 bp AMD-associated region

The AMD-associated 5,196 bp interval may harbor either unannotated genes or functional regulatory variants. To explore the latter possibility, we examined SNPs that could alter protein– DNA interactions within this region. Minor allele frequency (MAF) is the frequency of the less common allele in a given population. Analysis of 1000 Genomes Project data identified 229 SNPs spanning rs61871745 (Chr10:124,210,369) to rs3750846 (Chr10:124,215,565), a region in near-complete linkage disequilibrium with the established risk SNP rs10490924 (p.A69S; MAF = 0.273). Because SNPs in strong LD typically exhibit similar allele frequencies [30], we prioritized common variants (MAF > 0.1), reducing the list to 25 SNPs plus two flanking variants (27 total). Thirteen of these were previously reported by Grassmann et al. [22], whereas 12 were newly identified in our analysis (Figure 1A).

**Figure 1.**
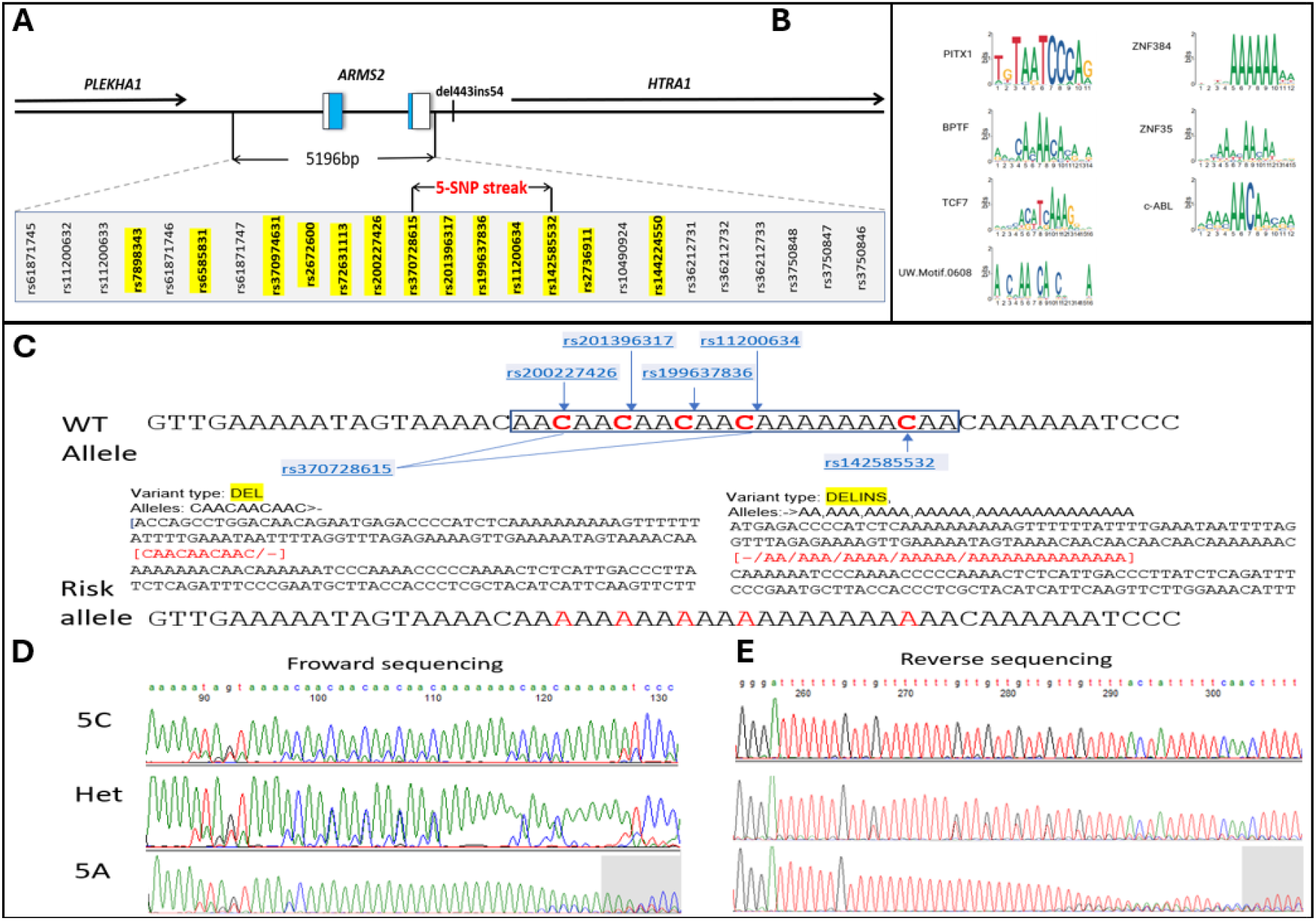
Consensus sequence of top 7 hits and 5 neighboring-SNP Streak sequences. **(A)** DNA sequence highlighting SNPs found in NCBI website are listed. **(B)** Sequence logos of 7 motifs of binding proteins. Sequence logo is a graphical representation of the DNA binding motif of a transcription factor/protein, based on the information content of each position. The information content ranges from 0, which corresponds to no consensus at the position, to 2, which corresponds to 1 consensus base used at the position. (**C)** Rs370728615 and rs142585532 were previously marked as deletion-insertion (DELINS) by NCBI. The figure indicates that all SNPs are C/A SNPs. **(D, E)** Forward and reverse Sanger sequencing reads of homozygous 5C (AA**<u>C</u>**AA**<u>C</u>**AA**<u>C</u>**AA**<u>C</u>**AAAAAAA**<u>C)</u>**, homozygous 5A Streaks (AA**<u>A</u>**AA**<u>A</u>**AA**<u>A</u>**AA**<u>A</u>**AAAAAAA**<u>A</u>) and** heterozygous (Het, 5C/5A)

Integrating motif scanning with binding free-energy prediction, our bioinformatic screen identified seven candidate trans-acting proteins—primarily transcription factors and kinases— whose binding is likely modulated by SNP variation (Figure 1B). Among these, c-ABL (cellular form of Abelson non-receptor tyrosine kinase) emerged as the top candidate, supported by both altered binding free energy and recurrent motif enrichment (Table 1, Table S1 and S2). The consensus c-ABL binding motif (5′-AA/CAACAAA/C-3′ [31]) was densely concentrated within a 28 bp segment (Chr10:122,454,671–122,454,688), located ∼550 bp upstream of the *ARMS2* start codon. This region contains multiple tandem motifs (AACAACAACAACAACAAAAAAACAACAA) overlapping five consecutive SNPs (highlighted in blue), suggesting that allelic variation within this “5-SNP streak” may disrupt c-ABL binding and thereby modulate transcriptional regulation relevant to AMD.

**Table 1.**
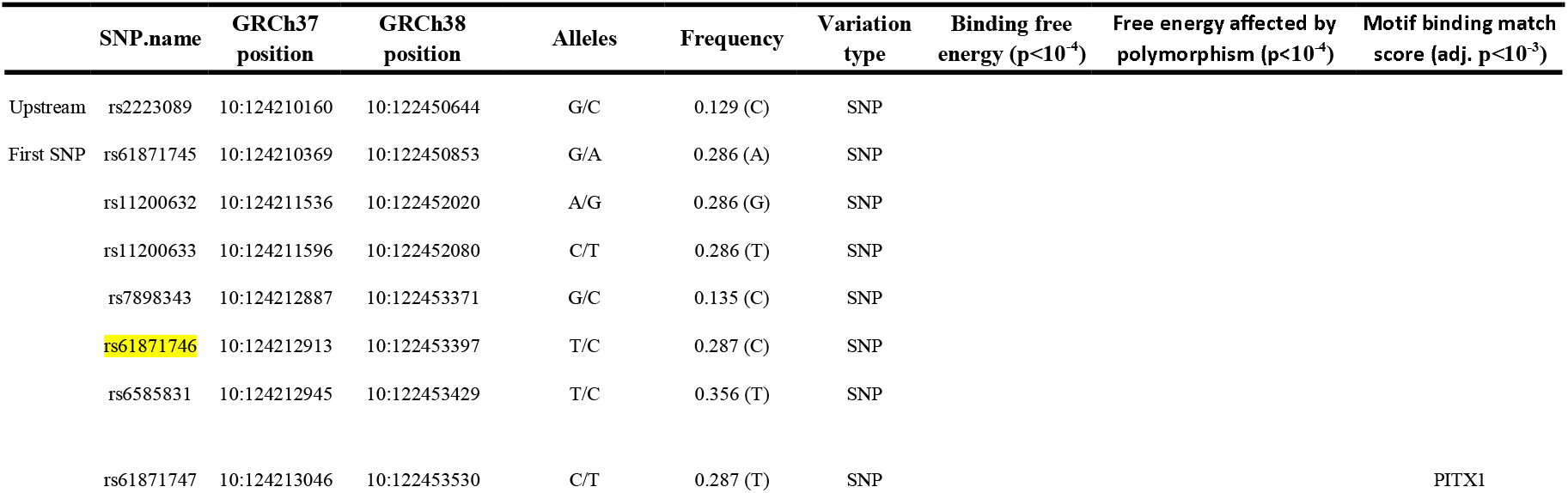
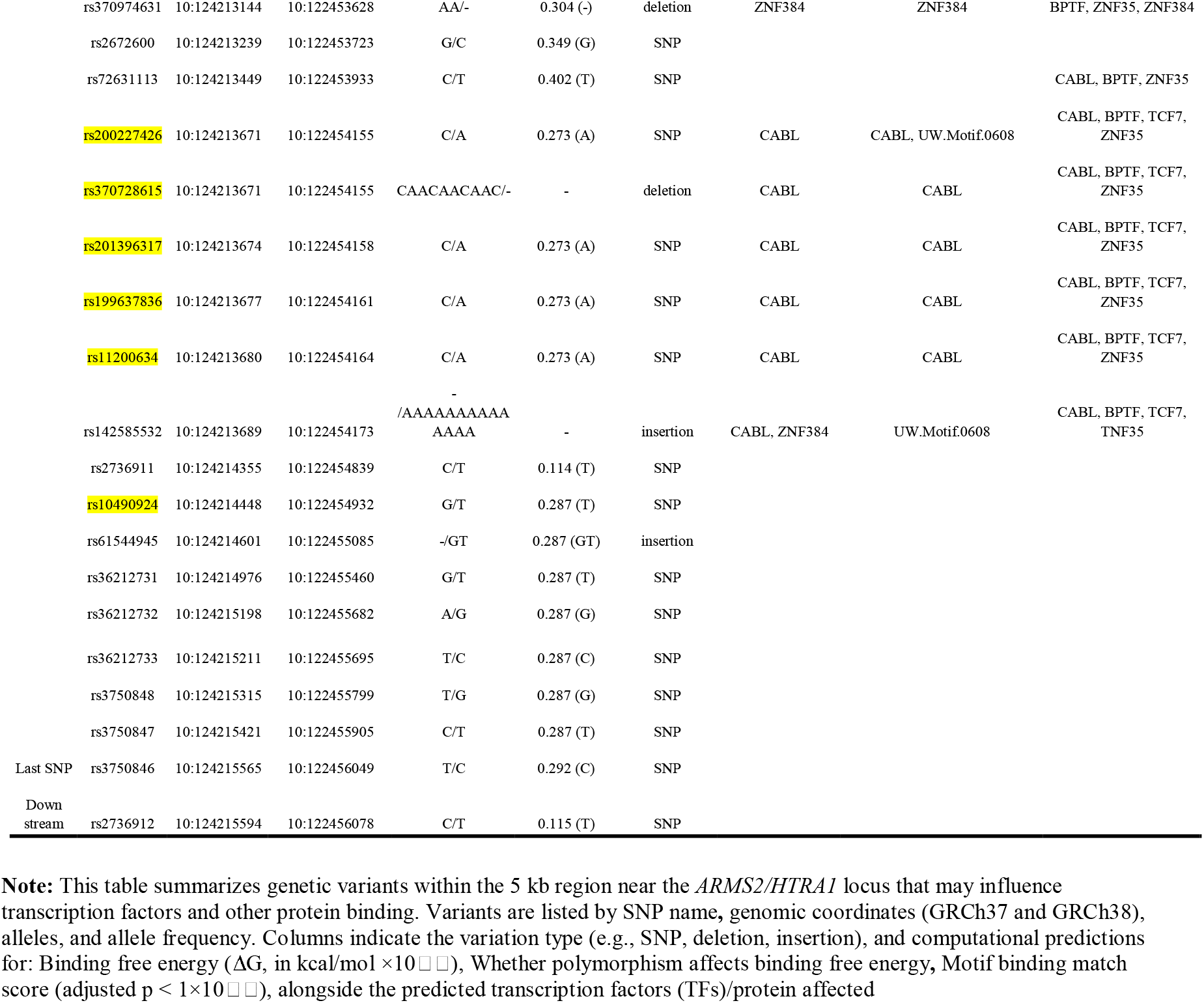
Putative regulatory candidates affecting protein binding within the 5,196-bp AMD-associated region.

|  | SNP.name | GRCh37 position | GRCh38 position | Alleles | Frequency | Variation type | Binding free energy (p<10<sup>-4</sup>) | Free energy affected by polymorphism (p<10<sup>-4</sup>) | Motif binding match score (adj. p<10<sup>-3</sup>) |
|---|---|---|---|---|---|---|---|---|---|
| Upstream | rs2223089 | 10:124210160 | 10:122450644 | G/C | 0.129 (C) | SNP |  |  |  |
| First SNP | rs61871745 | 10:124210369 | 10:122450853 | G/A | 0.286 (A) | SNP |  |  |  |
|  | rs11200632 | 10:124211536 | 10:122452020 | A/G | 0.286 (G) | SNP |  |  |  |
|  | rs11200633 | 10:124211596 | 10:122452080 | C/T | 0.286 (T) | SNP |  |  |  |
|  | rs7898343 | 10:124212887 | 10:122453371 | G/C | 0.135 (C) | SNP |  |  |  |
|  | rs61871746 | 10:124212913 | 10:122453397 | T/C | 0.287 (C) | SNP |  |  |  |
|  | rs6585831 | 10:124212945 | 10:122453429 | T/C | 0.356 (T) | SNP |  |  |  |
|  | rs61871747 | 10:124213046 | 10:122453530 | C/T | 0.287 (T) | SNP |  |  | PITX1 |
|  | rs370974631 | 10:124213144 | 10:122453628 | AA/- | 0.304 (-) | deletion | ZNF384 | ZNF384 | BPTF, ZNF35, ZNF384 |
|  | rs2672600 | 10:124213239 | 10:122453723 | G/C | 0.349 (G) | SNP |  |  |  |
|  | rs72631113 | 10:124213449 | 10:122453933 | C/T | 0.402 (T) | SNP |  |  | CABL, BPTF, ZNF35 |
|  | rs200227426 | 10:124213671 | 10:122454155 | C/A | 0.273 (A) | SNP | CABL | CABL, UW.Motif.0608 | CABL, BPTF, TCF7, ZNF35 |
|  | rs370728615 | 10:124213671 | 10:122454155 | CAACAACAAC/- | - | deletion | CABL | CABL | CABL, BPTF, TCF7, ZNF35 |
|  | rs201396317 | 10:124213674 | 10:122454158 | C/A | 0.273 (A) | SNP | CABL | CABL | CABL, BPTF, TCF7, ZNF35 |
|  | rs199637836 | 10:124213677 | 10:122454161 | C/A | 0.273 (A) | SNP | CABL | CABL | CABL, BPTF, TCF7, ZNF35 |
|  | rs11200634 | 10:124213680 | 10:122454164 | C/A | 0.273 (A) | SNP | CABL | CABL | CABL, BPTF, TCF7, ZNF35 |
|  | rs142585532 | 10:124213689 | 10:122454173 | -/AAAAAAAAAAAAAA | - | insertion | CABL, ZNF384 | UW.Motif.0608 | CABL, BPTF, TCF7, TNF35 |
|  | rs2736911 | 10:124214355 | 10:122454839 | C/T | 0.114 (T) | SNP |  |  |  |
|  | rs10490924 | 10:124214448 | 10:122454932 | G/T | 0.287 (T) | SNP |  |  |  |
|  | rs61544945 | 10:124214601 | 10:122455085 | -/GT | 0.287 (GT) | insertion |  |  |  |
|  | rs36212731 | 10:124214976 | 10:122455460 | G/T | 0.287 (T) | SNP |  |  |  |
|  | rs36212732 | 10:124215198 | 10:122455682 | A/G | 0.287 (G) | SNP |  |  |  |
|  | rs36212733 | 10:124215211 | 10:122455695 | T/C | 0.287 (C) | SNP |  |  |  |
|  | rs3750848 | 10:124215315 | 10:122455799 | T/G | 0.287 (G) | SNP |  |  |  |
|  | rs3750847 | 10:124215421 | 10:122455905 | C/T | 0.287 (T) | SNP |  |  |  |
| Last SNP | rs3750846 | 10:124215565 | 10:122456049 | T/C | 0.292 (C) | SNP |  |  |  |
| Down stream | rs2736912 | 10:124215594 | 10:122456078 | C/T | 0.115 (T) | SNP |  |  |  |

**Note:** This table summarizes genetic variants within the 5 kb region near the *ARMS2/HTRA1* locus that may influence transcription factors and other protein binding. Variants are listed by SNP name**,** genomic coordinates (GRCh37 and GRCh38), alleles, and allele frequency. Columns indicate the variation type (e.g., SNP, deletion, insertion), and computational predictions for: Binding free energy (ΔG, in kcal/mol ×10[illegible]), Whether polymorphism affects binding free energy**,** Motif binding match score (adjusted p < 1×10[illegible]), alongside the predicted transcription factors (TFs)/protein affected

Consistently, c-ABL was the most frequently predicted binding factor whose interaction was altered by sequence variation. SNPs within the previously defined 5-SNP AMD risk SNP streak (consecutive SNPs) showed the greatest predicted impact on c-ABL binding. Variants predicted to disrupt other motifs— including PITX1, BPTF, TCF7, ZNF35, and ZNF384—were also identified, supporting a broader regulatory impact of these polymorphisms on local chromatin and gene expression. C-ABL is the candidate for a top trans-acting factor whose binding may be affected by SNP polymorphism in the 5k region.

### Characterization of a functional 5-SNP streak in the *ARMS2* promoter region

The c-ABL-binding segment contained six consecutive SNPs, according to NCBI annotations. Four SNPs (rs200227426, rs201396317, rs199637836, rs11200634) shared an identical MAF of 0.273 in the 1000 Genomes Project, while rs370728615 and rs142585532 lacked reliable MAF data. To clarify variant status, we performed Sanger sequencing in 78 human DNA samples (24 controls, 23 wet AMD, 31 dry AMD). This analysis differed from NCBI annotations in that rs370728615 and rs142585532 appear as C/A SNPs rather than as deletion–insertion variants (Figure 1C).

Bidirectional sequencing was necessary to overcome limited read quality in the C-rich downstream flanking region (Figure 1D). These experiments confirmed the presence of five tightly linked SNPs -rs200227426, rs201396317, rs199637836, rs11200634, and rs142585532 -consistently forming a haplotype streak. This 5-SNP streak existed either as a major allele frequency allele (5C: AACAACAACAACAAAAAAACAA) or as a MAF allele (5A: AAAAAAAAAAAAAAAAAAAAAA). Four of these SNPs had identical allele frequencies in 1000 Genomes, consistent with their strong LD. Together, these results defined a 5-SNP regulatory streak/haplotype in the *ARMS2* promoter region.

### The 5-SNP streak is strongly associated with AMD

The 5-SNP streak we identified is not included among the 13 SNPs previously associated with AMD in the 5,196 bp region. Although this upstream region contains several previous reported SNPs in strong linkage disequilibrium (LD) with rs10490924, this does not imply that all SNPs in the region share high LD with it.

We next evaluated whether the 5-SNP streak is associated with AMD susceptibility. Genotyping in 438 AMD patients (179 dry AMD, 251 wet AMD) and 136 controls revealed that the 5A allele in the streak was significantly associated with both dry AMD (OR = 3.37; 95% CI: 2.35–4.84; p = 1.87 × 10□^11^) and wet AMD (OR = 3.80; 95% CI: 2.70–5.35; p = 3.28 × 10□^1^□) (Table 2).

**Table 2.** The 5-SNP Streak is significantly associated with both wet and dry forms of age-related macular degeneration (AMD)

| 5-SNP Streak | Non-AMD | Wet AMD |  |  | Dry AMD |  |  |
|---|---|---|---|---|---|---|---|
| Genotype | Control (%) | Cases (%) | OR (95% CI)<br>vs reference | P | Cases (%) | OR (95% CI<br>vs reference) | P |
| 5C homozygous | 89 (65.44) | 7 (31.8) |  |  | 63 (24.32) |  |  |
| 5C/5A heterozygous | 39 (28.68) | 8 (34.8) |  |  | 138 (53.28) |  |  |
| 5A homozygous | 8 (5.88) | 8 (34.8) |  |  | 58 (22.39) |  |  |
| 5C | 217 (79.78) | 22 (47.8) |  |  | 264 (50.96) |  |  |
| 5A | 55 (20.72) | 24 (52.2) | 3.37 (2.35-4.84) | 1.87e-11 | 254 (49.03) | 3.80 (2.70-5.35) | 3.28e-15 |
| Total alleles | 272 (100) | 358 (100) |  |  | 518 (100) |  |  |
| Total samples | 136 | 179 |  |  | 259 |  |  |

**Note:** Genotypic distribution of the 5C and 5A SNP haplotype Streaks in the non-AMD controls, wet AMD, and dry AMD patients. Genotypes include 5C homozygous, 5C/5A heterozygous, and 5A homozygous individuals. The table shows the number and percentage of individuals within each group, along with odds ratios (ORs), 95% confidence intervals (CIs), and p-values relative to the reference group (non-AMD)

Further analysis demonstrated that the risk allele 5A in the 5-SNP streak is in near-complete LD with the risk SNP T in the rs10490924 (r^2^ = 0.99; n = 574 samples), reinforcing its candidacy as a functional regulatory element and a surrogate genetic marker for AMD risk.

### Experimental validation confirms direct c-ABL binding to the 5-SNP streak

To assess the relevance of c-ABL in ocular tissues, we quantified its transcript levels across multiple human ocular samples. c-ABL mRNA was detected in human retina, choroid/RPE, iPSC-derived RPE, and ARPE-19 cells, with highest expression observed in choroid/RPE and iPSC-derived RPE relative to retina and ARPE-19 (Figure 2A).

**Figure 2.**
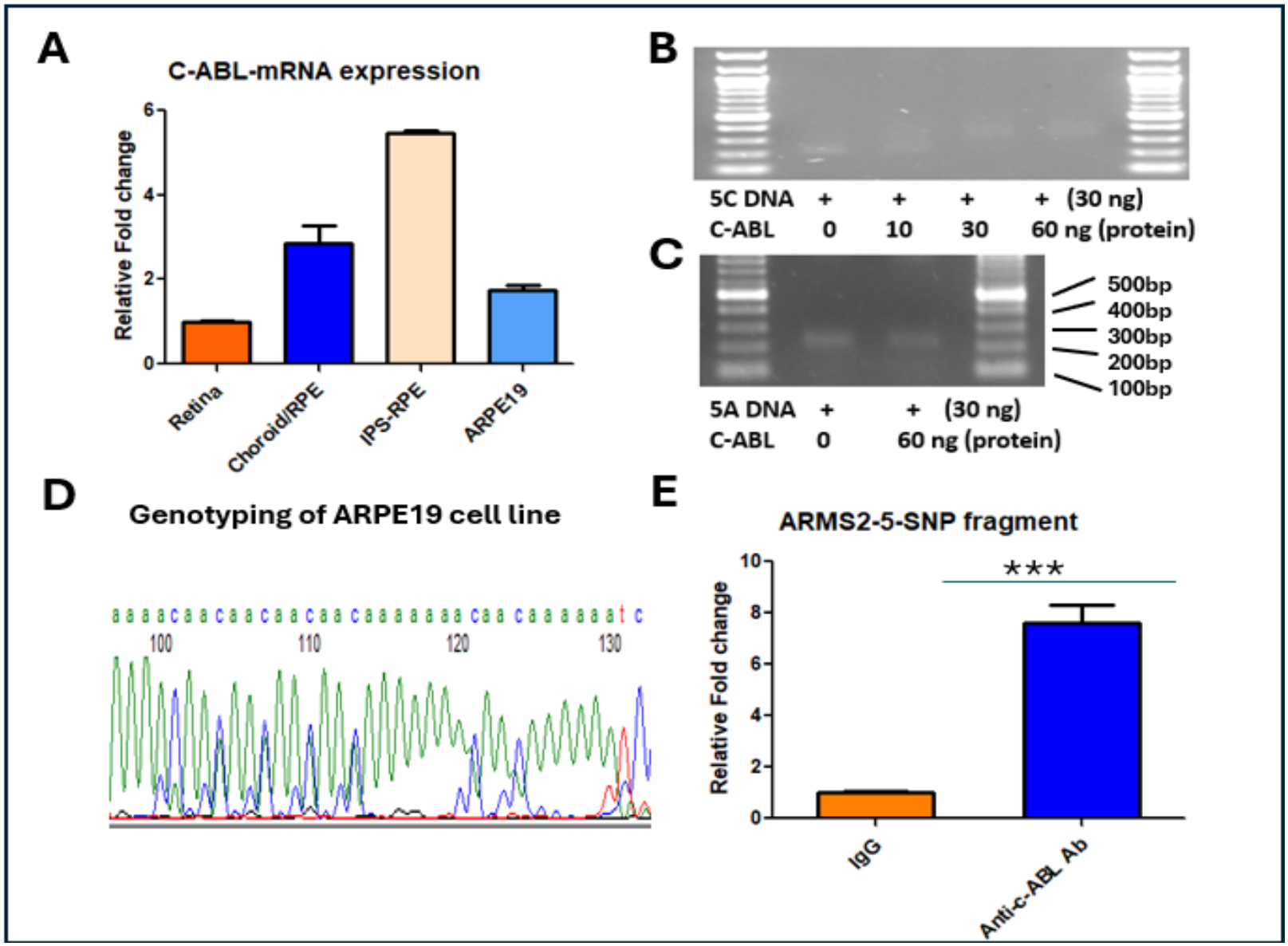
Verification of the binding of C-ABL transcription factor binding with the AMD-associated 5-SNP Streak. **(A)** Expression levels of C-ABL mRNA in different ocular cell types/tissues by qPCR, showing higher expression in iPSC-derived RPE and choroid/RPE compared to retina and ARPE19 cells. (**B, C)** EMSA shows c-ABL protein varied binding to the ARMS2-5-SNP fragment in human donor samples with different haplotypes: 5A (risk) and 5C (non-risk). **(D**) Representative sequencing trace showing genotyping of the ARPE19 cell line at the 5-SNP locus, confirming its heterozygous 5C/5A allelic configuration. **(E**) Quantification of ChIP in ARPE19 cells using an anti-C-ABL antibody vs. control IgG shows significant enrichment at the 5-SNP streak region

To determine whether c-ABL directly binds the AMD-associated region, we first performed electrophoretic mobility shift assays (EMSA) using recombinant c-ABL protein. A 220 bp DNA fragment carrying the non-risk 5C alleles exhibited a clear mobility shift relative to the risk 5A alleles, demonstrating allele-specific binding of c-ABL in vitro (Figure 2B-C).

Because EMSA assesses protein–DNA interactions in a cell-free environment, we next evaluated c-ABL binding in vivo using chromatin immunoprecipitation and quantitative PCR (ChIP-qPCR). ARPE-19 cells, confirmed by sequencing to be heterozygous at the 5-SNP streak (Figure 2D), were subjected to ChIP with an anti–c-ABL antibody. Quantitative PCR revealed a 7.63 ± 0.94– fold enrichment (P < 0.001) of heterozygous 5C/5A allele–containing DNA immunoprecipitated with the anti–c-ABL antibody compared with the IgG control, confirming in vivo c-ABL occupancy at this regulatory locus (Figure 2E).

### c-ABL binding represses transcriptional activity through the 5-SNP streak

To determine functional consequences of c-ABL binding to the 5-SNP streak, we cloned a 2.3 kb *ARMS2* upstream fragment containing either the 5C (non-risk) or 5A (risk) allele into luciferase reporter constructs (Figure 3A). Baseline reporter assays in HEK293T and ARPE-19 cells showed no significant differences between the 5C and 5A alleles (Figure 3B–C), consistent with previous findings [32].

**Figure 3.**
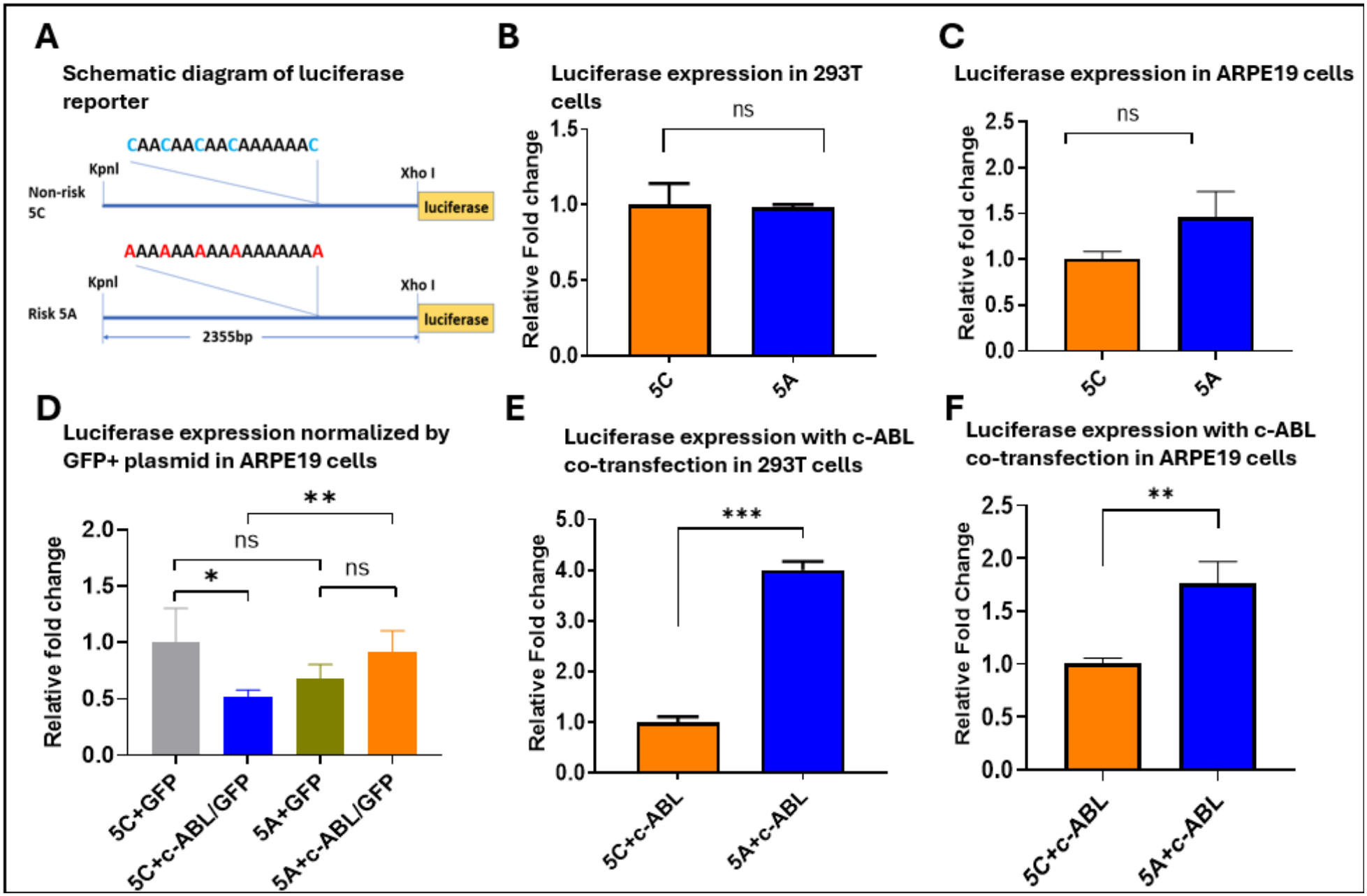
Functional analysis of the non-risk 5C and risk 5A alleles within the 5-SNP streak. (A) Schematic representation of luciferase reporter constructs containing either the non-risk 5C or risk 5A SNP streak. The 5C allele harbors the canonical sequence “AACAACAACAACAAAAAAACAA,” whereas the risk 5A allele contains an uninterrupted poly(A) tract (“AAAAAAAAAAAAAAAAAAAAAA”). A 2,355-bp fragment encompassing the SNP streak was cloned upstream of the luciferase reporter using KpnI and XhoI restriction sites. (B, C) Luciferase reporter activity in HEK293T and ARPE-19 cells transfected with the 5C or 5A constructs. No significant difference in basal transcriptional activity was observed between the two alleles. (D) Luciferase activity, normalized to a GFP-positive plasmid, in ARPE-19 cells co-transfected with c-ABL and either the 5C or 5A construct. (E, F) Luciferase activity, normalized to Renilla luciferase, in HEK293T and ARPE-19 cells co-transfected with c-ABL and either the 5C or 5A reporter construct

However, allele-specific effects became evident upon co-transfection with a c-ABL expression plasmid. c-ABL overexpression markedly suppressed luciferase activity driven by the wild-type 5C construct, whereas the mutant 5A construct was largely resistant to this repression (Figure 3D). Quantitatively, luciferase activity from the 5A construct was increased by approximately 4.00 ± 0.24-fold in HEK293T cells and 1.75 ± 0.26-fold in ARPE-19 cells relative to the 5C construct (Figure 3E-F). These findings indicate that c-ABL binding to the 5C allele mediates transcriptional repression, while the 5A allele disrupts this regulatory interaction and abrogates c-ABL-dependent repression.

### *ARMS2* mRNA expression increased in 5A genotype compared with 5C in human RPE/choroid tissues

To determine whether A*RMS2* mRNA expression differs between non-risk and risk haplotypes, we quantified *ARMS2* transcript levels in human RPE/choroid samples carrying homozygous non-risk 5C (n=3), heterozygous 5C/5A (n=4), or homozygous risk 5A (n=3) genotypes of the 5-SNP haplotype. *ARMS2* mRNA abundance was lowest in samples homozygous for the 5C genotype. In contrast, samples homozygous for the 5A risk haplotype exhibited a significant increase in *ARMS2* expression, reaching 1.73 ± 0.06-fold relative to 5C samples (P < 0.001). Heterozygous 5C/5A samples did not show a significant increase compared with homozygous 5C samples (Figure 4A). In contrast, *HTRA1* mRNA expression did not differ significantly among the three genotypes (Figure 4B).

**Figure 4.**
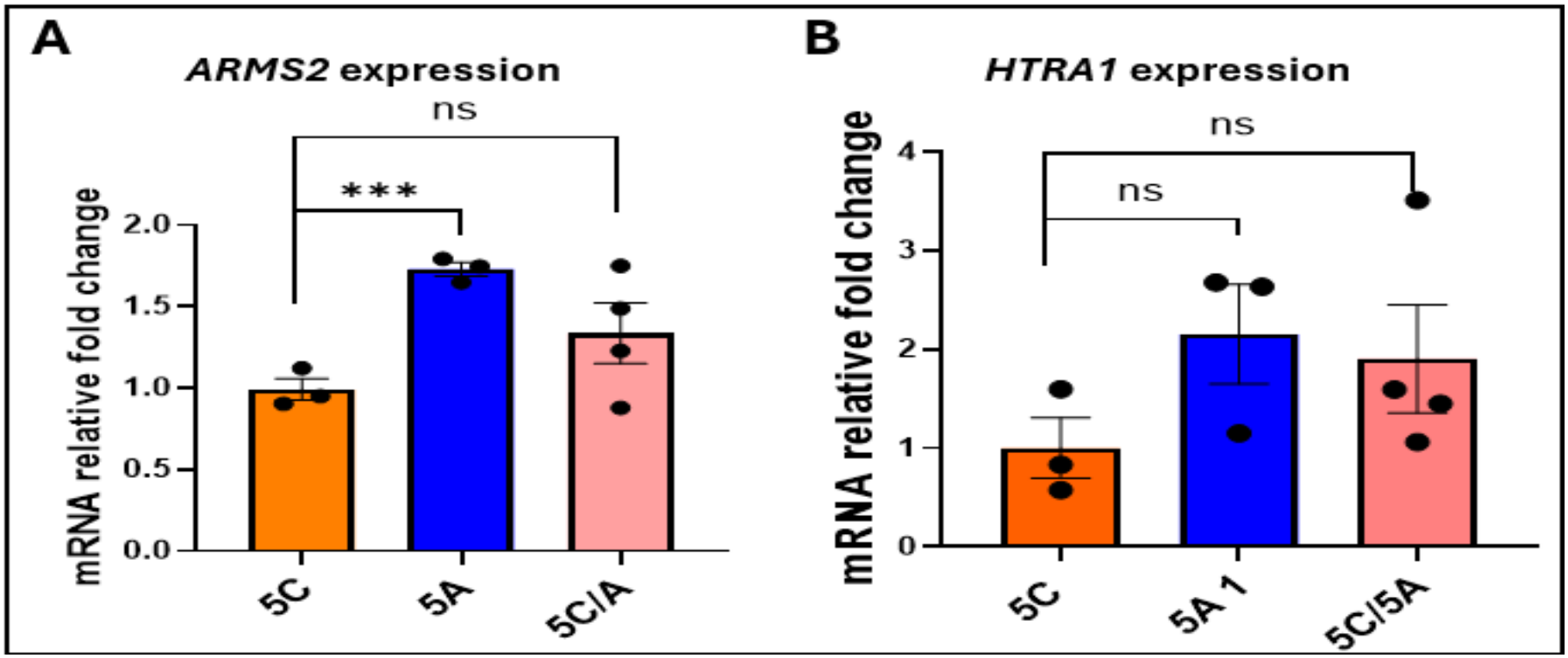
*ARMS2* and *HTRA1* mRNA expression in human RPE/choroid samples stratified by genotype. (A) Relative *ARMS2* mRNA expression and (B) relative *HTRA1* mRNA expression measured in human RPE/choroid tissue. Samples are grouped by genotypes (5C, 5A, and 5C/5A). Bars represent mean ± SEM, and individual dots denote independent biological samples. Statistical significance was assessed using Student’s t-test with appropriate multiple comparisons

Together, these results indicate that the AMD-associated 5A haplotype is specifically associated with elevated *ARMS2* mRNA expression in human RPE/choroid tissue, supporting a genotype-dependent regulatory effect.

## Discussion

The 10q26 risk haplotype has consistently emerged as one of the strongest genetic signals for AMD, refining the susceptibility interval to the ARMS2–HTRA1 locus. This has prompted speculation that ARMS2 may represent the causal gene or that critical regulatory elements reside within its promoter. However, genetic association alone cannot establish causality, and the precise variants driving AMD risk remain unresolved. Most variants in this region, including the well-studied missense SNP rs10490924 (p.A69S), have uncertain functional consequences, and their pathogenic relevance continues to be debated [10].

Previous investigations have primarily focused on cis-regulatory mechanisms and protein-coding functions of ARMS2 and HTRA1. Although cis-effects within the 5,196 bp promoter region have been reported, potential trans-regulatory influences have been largely overlooked. Given the density of associated SNPs—25 common variants within this short interval—it is plausible that multiple variants act in concert to influence gene regulation, keeping both ARMS2 and HTRA1 as viable candidates for AMD pathogenesis.

In this study, we identified c-ABL as the strongest candidate trans-acting protein interacting with a 5-SNP block in the ARMS2 promoter region. Originally characterized as the mammalian homolog of the Abelson murine leukemia virus oncogene [33], c-ABL is a non-receptor tyrosine kinase with key roles in DNA damage responses, oxidative stress signaling, and neuronal cell death [34]. Our in vitro analyses demonstrated that this noncoding 5-SNP streak can modulate transcriptional activity in a c-ABL–dependent manner, supporting a model in which sequence variation disrupts repressor binding and thereby alters gene expression.

The observed increase in ARMS2 mRNA levels in RPE/choroid samples carrying the AMD-associated 5A SNP streak haplotype provides additional evidence that genetic variation at the ARMS2/HTRA1 locus influences transcriptional output in ocular tissues. Elevated ARMS2 expression in 5A donors is consistent with prior reports linking the risk haplotype to altered regulatory activity and supports a model in which noncoding variants modulate ARMS2 transcription through changes in transcription factor binding, potentially including c-ABL, as suggested by our EMSA and ChIP analyses. Given ARMS2’s roles in mitochondrial maintenance and extracellular matrix dynamics, upregulation associated with the 5A risk genotype may contribute to RPE dysfunction or increased susceptibility to stress in AMD. These findings reinforce the functional relevance of the risk haplotype and highlight ARMS2 dysregulation as a plausible mechanistic contributor to disease pathogenesis.

Conflicting results have been reported regarding ARMS2 expression. One study suggested that the AMD risk haplotype reduces ARMS2 mRNA abundance, potentially via destabilizing mutations in the RPE or mechanisms that remain unclear in the retina [35]. In contrast, another study reported that AMD-associated variants at chromosome 10q26 do not significantly affect ARMS2 or HTRA1 transcript levels in the human retina [36]. A key limitation of both these studies, as well as ours, is the overall low expression of ARMS2 in RPE and retina, reflected by qPCR Ct values approaching 34. Gene expression profiling studies of these tissues similarly report very low baseline ARMS2 expression, making detection of differential expression challenging without large sample sizes [37]. An additional limitation of our study is the small sample size, with only 3–4 samples per genotype available for comparison.

Our findings expand current understanding of the 10q26 locus by implicating a trans-acting regulatory mechanism in addition to previously reported cis effects. Nonetheless, limitations remain. Our binding screen was restricted to transcription factors and kinases catalogued in existing motif databases, and additional relevant regulators may have been missed. Moreover, while our assays demonstrate binding and functional effects, the downstream gene targets and tissue-specific consequences of altered c-ABL binding remain to be fully elucidated.

In summary, we define a novel 5-SNP regulatory streak upstream of ARMS2 that directly binds c-ABL and influences transcriptional regulation. This discovery highlights noncoding elements as potential drivers of AMD risk and provides a framework for future studies aimed at pinpointing causal variants and mechanistic pathways. Ultimately, resolving the functional impact of these variants may uncover new molecular targets and therapeutic strategies for AMD.

## Contributors

PWZ initiated the idea, designed all the experiments and wrote the main manuscript text. SL did the Bioinformatic screening for trans-acting elements binding; WL, LF, SL, ZHW, JA and CAB made significant contribution to the experiment performance, data collection and analysis. PWZ, SLM and DJZ supervised the whole process of data collection, analysis, results interpretation and manuscript writing.

## Declaration of Interests

The authors declare no conflicts of interests.

## Data sharing

The authors declare that all data generated or analysed during this study are included in this published article.

## Acknowledgement

The present study was supported by NEI P30 EY001765 (Wilmer Core Grant, Microscopy Module).

## Notes

### Competing Interest Statement

The authors have declared no competing interest.

### Summary of Updates

We have revised the manuscript to improve its clarity, scientific rigor, and presentation of the findings. The major revisions are summarized below: Bioinformatic and genetic analyses: Clarified the identification of the 5-SNP cluster within the ARMS2 promoter region and its association with AMD susceptibility, including its strong linkage disequilibrium with rs10490924. c-ABL binding characterization: Improved the presentation of EMSA and ChIP results supporting allele-dependent c-ABL binding to the 5-SNP regulatory sequence. Transcriptional regulation: Refined the interpretation of luciferase reporter assays demonstrating allele-specific transcriptional regulatory effects. Discussion and presentation: Strengthened the discussion of the potential molecular mechanisms linking the 5-SNP cluster to AMD and improved the clarity of the text, figures, and conclusions. Overall, these revisions strengthen the evidence supporting a potential role for the 5-SNP cluster in AMD susceptibility through altered c-ABL binding and transcriptional regulation.

