## supplementary file for "A Bioinformatically Identified 5-SNP Streak in the ARMS2 Promoter Region Links to AMD Through Altered c-ABL Binding and Transcriptional Regulation"

**Supplementary data**

**Table S1** **Counts and Frequencies of Trans-Acting Regulatory Element Binding at SNPs within the 5,196 bp segment encompassing the ARMS2 promoter**

| **Gene** | **Count** | **Frequency** |
| --- | --- | --- |
| CABL | 8 | 0.027491409 |
| FOXC1 | 6 | 0.020618557 |
| FOXC2 | 6 | 0.020618557 |
| FOXP1 | 6 | 0.020618557 |
| RREB1 | 6 | 0.020618557 |
| MEF2A | 5 | 0.017182131 |
| FOXP2 | 4 | 0.013745704 |
| HNF1B | 4 | 0.013745704 |
| IRF1 | 4 | 0.013745704 |
| MEF2C | 4 | 0.013745704 |
| CDX1 | 3 | 0.010309278 |
| FIGLA | 3 | 0.010309278 |
| FOSL2 | 3 | 0.010309278 |
| FOXL1 | 3 | 0.010309278 |
| HOXC10 | 3 | 0.010309278 |
| HOXD11 | 3 | 0.010309278 |
| LMX1B | 3 | 0.010309278 |
| MEF2D | 3 | 0.010309278 |
| ONECUT3 | 3 | 0.010309278 |
| PHOX2A | 3 | 0.010309278 |
| TCF7L2 | 3 | 0.010309278 |
| YY1 | 3 | 0.010309278 |
| BATF::JUN | 2 | 0.006872852 |
| BSX | 2 | 0.006872852 |
| CDX2 | 2 | 0.006872852 |
| DLX6 | 2 | 0.006872852 |
| ESX1 | 2 | 0.006872852 |
| EVX1 | 2 | 0.006872852 |
| FOS | 2 | 0.006872852 |
| FOS::JUN | 2 | 0.006872852 |
| FOXB1 | 2 | 0.006872852 |
| FOXI1 | 2 | 0.006872852 |
| GBX1 | 2 | 0.006872852 |
| GBX2 | 2 | 0.006872852 |
| HESX1 | 2 | 0.006872852 |
| HOXA5 | 2 | 0.006872852 |
| HOXC11 | 2 | 0.006872852 |
| HOXC12 | 2 | 0.006872852 |
| LEF1 | 2 | 0.006872852 |
| LHX9 | 2 | 0.006872852 |
| LMX1A | 2 | 0.006872852 |
| MEF2B | 2 | 0.006872852 |
| MIXL1 | 2 | 0.006872852 |
| MLXIPL | 2 | 0.006872852 |
| MSX1 | 2 | 0.006872852 |
| MSX2 | 2 | 0.006872852 |
| NFATC3 | 2 | 0.006872852 |
| NKX2-3 | 2 | 0.006872852 |
| ONECUT2 | 2 | 0.006872852 |
| PAX4 | 2 | 0.006872852 |
| POU2F2 | 2 | 0.006872852 |
| POU4F2 | 2 | 0.006872852 |
| PROP1 | 2 | 0.006872852 |
| PRRX1 | 2 | 0.006872852 |
| RHOXF1 | 2 | 0.006872852 |
| SHOX | 2 | 0.006872852 |
| SNAI2 | 2 | 0.006872852 |
| SP1 | 2 | 0.006872852 |
| SPI1 | 2 | 0.006872852 |
| SPIB | 2 | 0.006872852 |
| TBX15 | 2 | 0.006872852 |
| TBX5 | 2 | 0.006872852 |
| TCF3 | 2 | 0.006872852 |
| TCF4 | 2 | 0.006872852 |
| TFAP4 | 2 | 0.006872852 |
| UNCX | 2 | 0.006872852 |
| USF1 | 2 | 0.006872852 |

Note: A list of transcription factors predicted to bind the 5-SNP block region upstream of the ARMS2 gene. The table shows the number of predicted binding events (**Count**) and their relative **Frequency** across all identified binding instances. CABL was the most frequently predicted binder, followed by FOXC1, FOXC2, FOXP1, and RREB1.

**Table S2 Trans-Acting Regulatory Elements and Their Predicted Binding Sites**

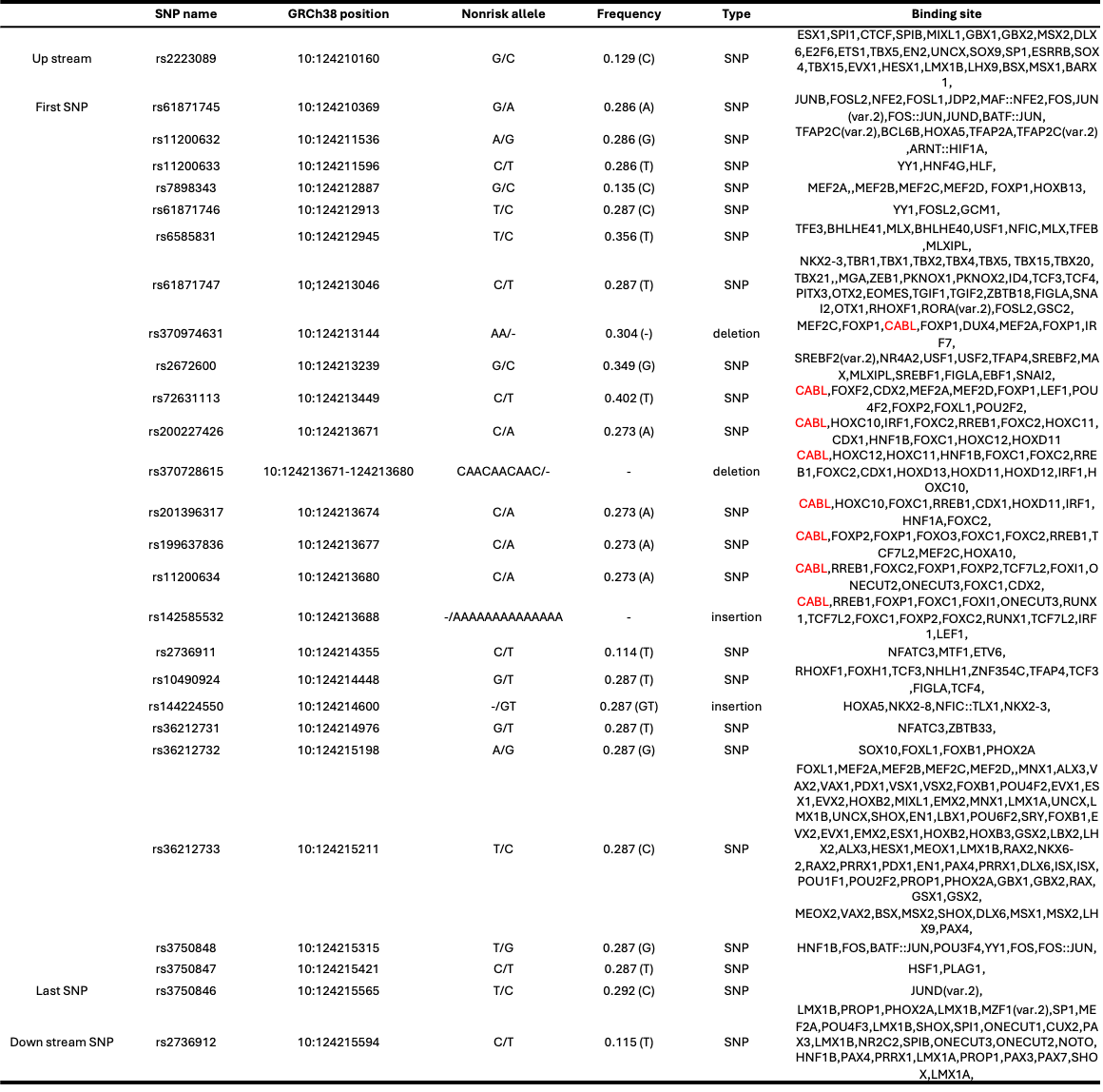

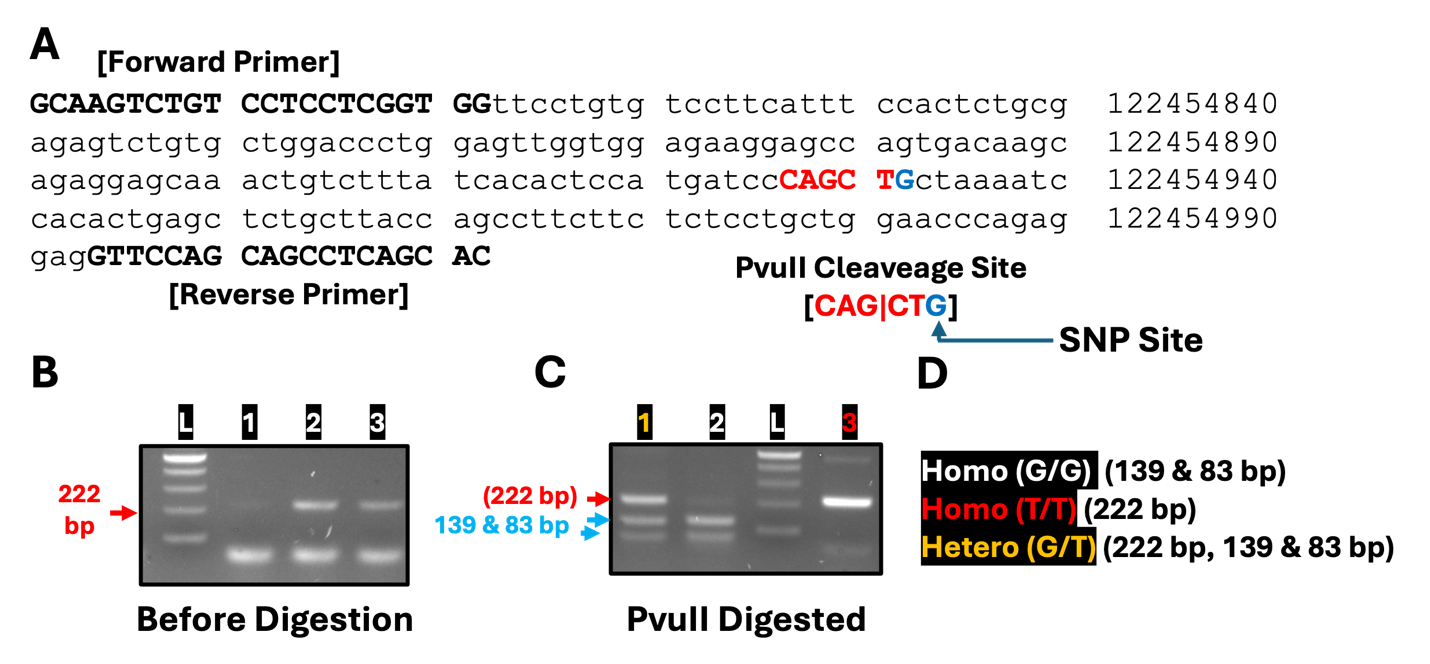

**Figure S1. Genotyping of the SNP rs10490924 (A69S) in the ARMS2 promoter region.** (A) Schematic of the SNP site showing the sequence context. The non-risk allele (G) is indicated in blue, respectively. The binding motif recognized by the trans-acting factor is highlighted above the sequence. The arrow indicates the SNP position within the motif. (B) Representative PCR-RFLP analysis of samples. Lane L, DNA ladder; lanes 1–3, individual samples. The 222 bp band corresponds to the undigested fragment. (C-D) Summary of genotype assignment after PvuII digestion: Homozygous G/G (white), Homozygous T/T (red), and Heterozygous G/T (yellow). Digestion of the homo G/G allele produces 139 and 83 bp fragments (Homo G/G). Digestion of the heterozygous G/T produces 139, 83 bp and 222 bp fragments; Digestion of the homo T/T produces 222bp band (no cut for pvuII restriction enzyme
